# Gestational Diabetes Mellitus Is a Risk for Macrosomia: Case- Control Study in Eastern Ethiopia

**DOI:** 10.1101/492355

**Authors:** Elias Bekele Wakwoya, Tariku Dingeta Amante, Kasahun Fikadu Tesema

## Abstract

**Background:** Gestational diabetes mellitus is any degree of glucose intolerance at onset or first recognition during pregnancy. A pregnant woman with diabetes and her unborn child are at increased risk of pregnancy complications and adverse neonatal outcomes. The aim of this study was to assess the association of gestational diabetes mellitus and adverse birth outcomes among women who gave birth in Eastern Ethiopia.

**Method:** Unmatched case control study design was conducted in Hiwot Fana Specialized University Hospital and Dilchora Hospital from December 2015 to April 2017. This study involved a total of 1,834 mothers and their babies. A structured and pre-tested questionnaire was used to collect the socio-demographic data. Mothers who had risk factor for gestational diabetes were screened by oral glucose tolerance tests. Adverse birth outcomes were observed and registered after delivery. Multivariate logistic regression analysis was employed to identify predictors of adverse birth outcome. P value less than 0.05 was considered to decide statistical significance.

**Results:** From a total of 1,834 mothers 47 (2.6%) of them were found to have gestational diabetes. In binary logistic regression analysis macrosomia and still were found to have an association with gestational diabetes, COR=11[95% CI = 5.7-21.2] and COR= 2.9[95% CI = 1.02-8.5] respectively. Macrosomia was independently associated with GDM and babies born to mothers with gestational diabetes. Babies born from mothers with gestational diabetes were 8.5 times more likely to have macrosomia than babies born to non-diabetic mothers, AOR = 8.5 [95% CI = 5.7-21.4].

**Conclusion:** This study revealed that only macrosomia was strongly associated with gestational diabetes and this finding is coherent with studies done at different parts of the world. Early screening and treatment of mothers with GDM can minimize the adverse birth outcomes, therefore routine screening service for pregnant women who are at risk of developing gestational diabetes must exist at all health facilities in Ethiopia.

## Introduction

Diabetes mellitus is the common medical complication of pregnancy and it carries high risk to the fetus and the mother. Gestational diabetes mellitus is defined as being any degree of glucose intolerance at onset or first recognition during pregnancy and should include glucose readings that fall within the impaired glucose tolerance (IGT) diagnostic range, as well as those within the diagnostic range for diabetes [1,2]. The American Diabetes and Association also defines GDM as diabetes diagnosed during pregnancy that is not clearly overt diabetes [3].

The number of people with diabetes is increasing globally. As the incidence of diabetes continues to rise and increasingly affects individuals of all ages, pregnant women and their babies are at increased risk of diabetes [4]. The prevalence of gestational diabetes mellitus ranges from 2% - 14% of all pregnancies worldwide and 0 % - 13.9% in Africa [5, 6]. Even though there is no data for the total prevalence of GDM in Ethiopia, according to the study done in northern part of Ethiopia it was 3.7 percent [7].

Gestational diabetes can negatively affect the pregnancy and may be associated with many maternal, fetal and neonatal complications, both short and long term. A woman with GDM have an increased risk of adverse neonatal outcomes as compared to women without GDM. As different studies indicated, GDM is associated with a greater risk of neonatal fetal macrosomia, shoulder dystocia, neonatal trauma, respiratory distress, increased admission to neonatal intensive care units [8–12].

Adverse outcomes in pregnancies among women with diabetes are in most cases preventable by optimizing glycemic control. Early screening and treatment of mothers with GDM can minimize the complications for both mothers and their babies. Addressing GDM also constitutes a window of opportunity for early intervention and reduction of the future burden of type 2 diabetes [13]. However, in some of the poorest countries of the world, difficulties in accessing and receiving both maternity and general medical care increase the risks pregnant women to face complication of diabetes in pregnancy.

There was no information on the association of gestational diabetes with neonatal adverse neonatal outcome in our study area. Therefore, this study was conducted with the intention of assessing the association of GDM and adverse birth outcomes.

## Methods and materials

### Study area

This study was conducted in the labor wards of Hiwot Fana Specialized University Hospital and Dilchora Referral Hospital from December 2015 to April 2017. Hiwot Fana University Hospital is found in Harar city which is located 526km from Addis Ababa in the eastern part of Ethiopia. According to the Central Statistics Authority of Ethiopia in 2007, Harari regional state has population of 183,415 of which 92,316 were male and 91,099 were female [14]. Hiwot Fana Specialized University Hospital was established in 1941. It is referral hospital for both Harar town and its surroundings.

Dilchora Referral Hospital is found in Dire Dawa city administration council, located 501 km to the east of Addis Ababa. The hospital is serving an estimated population of 2 million, coming from Dire Dawa City administration and nearby Oromia and Somali regions. The hospital has a total number of 268 beds distributed between medical, pediatrics, surgical, gynecology and obstetrics wards. Monthly, an estimated 582 clients visit the antenatal clinic found in the hospital. In addition, an estimated 194 clients visit the clinic for antenatal care (ANC) each month.

### Study design and population

Institution based unmatched case control study design were conducted. The cases were women in labor who had gestational diabetes diagnosed after OGTT and controls were women who were not diagnosed with GDM. Mothers who were known diabetic before pregnancy and who were severely ill during data collection period were excluded from the study.

### Sample size determination and sampling procedure

Based on sample size determination for case control with assumption of prevalence of exposure among case, prevalence exposure among control and case to control ration of 1:4 minimum of 45 cases and 180 controls were needed. To obtain minimum number of cases involved in the study 1834 mother who came for delivery service were screened for gestational diabetes mellitus. To recruit study participants all mothers came to both hospitals during the study period were interviewed by using the structured questionnaire until the required sample size was obtained.

### Variables and measurement

Birth outcome were categorized as adverse birth outcomes if the babies had either of macrosomia, prematurity, admission to NICU, respiratory distress and congenital anomalies.

Pregnant women are considered as at risk for GDM if she had a history of delivery of previous macrosomic baby, history of stillbirth, family history of DM, obesity (BMI > 30kg/m^2^), previous congenital abnormal fetus and glucosuria. The babies were considered as macrosomic if their birth weight were greater than 4000gm.

Gestational Diabetes is any degree of glucose intolerance detected for the first time during pregnancy after 20 weeks of gestation and must fulfilled criteria listed by WHO (If RBS ≥140 mg/dL the patients will undergo a 3 hours 100 gm oral glucose tolerance test and GDM will be diagnosed if ≥ 2 values met or exceed the following cut-off point; blood sugar level at 1 hour - 190 mg/ dl, at 2 hours -165 mg/ dl, and at 3 hours - 145 mg/dl). OGTT is a provocation test to examine the efficiency of the body to metabolize glucose and it distinguishes metabolically healthy individuals from people with impaired glucose tolerance and those with diabetes. Random Blood Sugar is the amount of glucose dissolved in circulating blood, recorded irrespective of when food was last ingested.

### Data collection procedure

Socio-demographic data of the mothers were obtained on a face-to-face interviews. Screening for GDM were performed for mothers who were identified as at risk for developing GDM. For the mothers who had a RBS ≥140 mg/dl, 50 gm glucose challenge test were given orally to the mothers with the serum glucose measured one hour later. To ensure the quality of data a pretest was performed before actual data collection started and two days training was given for two days. All data were checked for completeness, clarity and consistency immediately after data collection.

### Data management and statistical analysis

Data were entered into EPI-info version 3.5.1 and then exported to SPSS version 20.0 software for analysis. After cleaning, frequencies and percentages were calculated. Bivariate and multivariate logistic regression analysis were used to ascertain any significant difference in any of adverse birth outcome among mother with GDM and without GDM. The adverse birth outcome include preterm birth, respiratory distress, macrosomia, stillbirth and congenital anomalies. P value of less than 0.05 was considered statistically significant.

### Ethical considerations

The protocol was approved by the Haramaya University Institutional Health Research Ethical Review Committee. Written and signed informed consent was obtained from each study participant prior to interview. Those mothers who were diagnosed with gestational diabetes were linked with the service where the treatment was given and all babies who had adverse birth outcomes were also managed accordingly.

## Results

### Demographic and obstetric characteristics

A total of 1834 women who came to Hiwot Fana and Dilchora Hospitals for delivery service were included in the study. The mean age of mothers for cases and controls was 25.6 ± 4.83 and 26.8 ± 4.59 respectively. The magnitude of married women were 42(89.4%) in cases and 1702(95.2%) in controls. From the total participants 29(61.7%) of cases and 1067(59.7%) of controls were educated. The proportion of mothers who had family history of DM was 34% in cases and 2.4% in controls. Among control group 644 (36%) and 15 (32%) of cases gave birth for the first time whereas 531 (30%) of control group and 8 (17%) of cases were pregnant for the first time. More than half of the cases 31(66%) and 215 (12.3%) of controls had pre-pregnancy obesity (Table 1).

**Table 1:** Socio demographic and selected obstetrics characteristics of women who gave birth in Hiwot fana and Dilchora hospitals from June 2016 to April 2017 G.C.

| Variables | Gestational diabetes |  |
| --- | --- | --- |
|  | Cases N (%) | Controls N (%) |
| **Age of mothers** |  |  |
| <18 | 0 (0%) | 21 (1.2%) |
| 18-34 | 2 (4.3%) | 33 (1.9%) |
| >35 | 45 (95.7%) | 1733 (96.9%) |
| **Marital status** |  |  |
| Single | 0 (0%) | 19 (1.1%) |
| Married | 42 (89.4%) | 1702 (95.2%) |
| Divorced | 4 (8.5%) | 38 (2.1%) |
| Widowed | 1 (2.1%) | 28 (1.6%) |
| **Ethnicity** |  |  |
| Oromo | 33 (70.2%) | 1171 (65.6%) |
| Amahara | 8 (17.0%) | 351 (19.6%) |
| Others | 6 (12.8%) | 265 (14.8%) |
| **Religion** |  |  |
| Christian | 15 (31.9%) | 544 (30.4%) |
| Muslim | 32 (68.1%) | 1243 (69.6%) |
| **Occupation** |  |  |
| Employee | 9 (19.1%) | 456 (25.5%) |
| Housewife | 32 (68.1%) | 1048 (58.7%) |
| Merchant | 6 (12.8%) | 283 (15.8%) |
| **Educational status** |  |  |
| Uneducated | 18 (38.3%) | 720 (40.3%) |
| Educated | 29 (61.7%) | 1067 (59.7%) |
| **Income** |  |  |
| < 1000 | 11 (23.4%) | 606 (33.9%) |
| > 1000 | 36 (76.6%) | 1181 (66.1%) |
| **Gravidity** |  |  |
| Primigravida | 8 (17.0%) | 531 (29.7%) |
| Multigravida | 39 (83.0%) | 1256 (70.3%) |
| **Parity** |  |  |
| Primipara | 15 (32.0%) | 644 (36.0%) |
| Multipara | 32 (68.0%) | 1143 (64.0%) |
| **Family history of DM** |  |  |
| No | 31(66.0%) | 1745 (94.5%) |
| Yes | 16 (34.0%) | 42 (5.5%) |

### Birth Outcomes

Preterm delivery among cases was 10.6% and 11.2% among controls. The proportion of malpresentation among cases and controls was 17% and 13.5% respectively. Stillbirth was presented in 8.5% of cases and 3.1% of controls. The proportion of macrosomia was higher among cases (31.9%) than controls (4.1%). From the total study participants 15% of neonates of cases were admitted to NICU while 10.7% of neonates of controls were admitted to NICU. Only 6.4% and 2.9% of neonates were born with congenital anomalies among cases and controls respectively. From the total participants, 8 (17%) of neonates among cases and 205 (11.5%) of neonates among controls had developed neonatal respiratory distress. (Fig 1).

**Fig 1:**
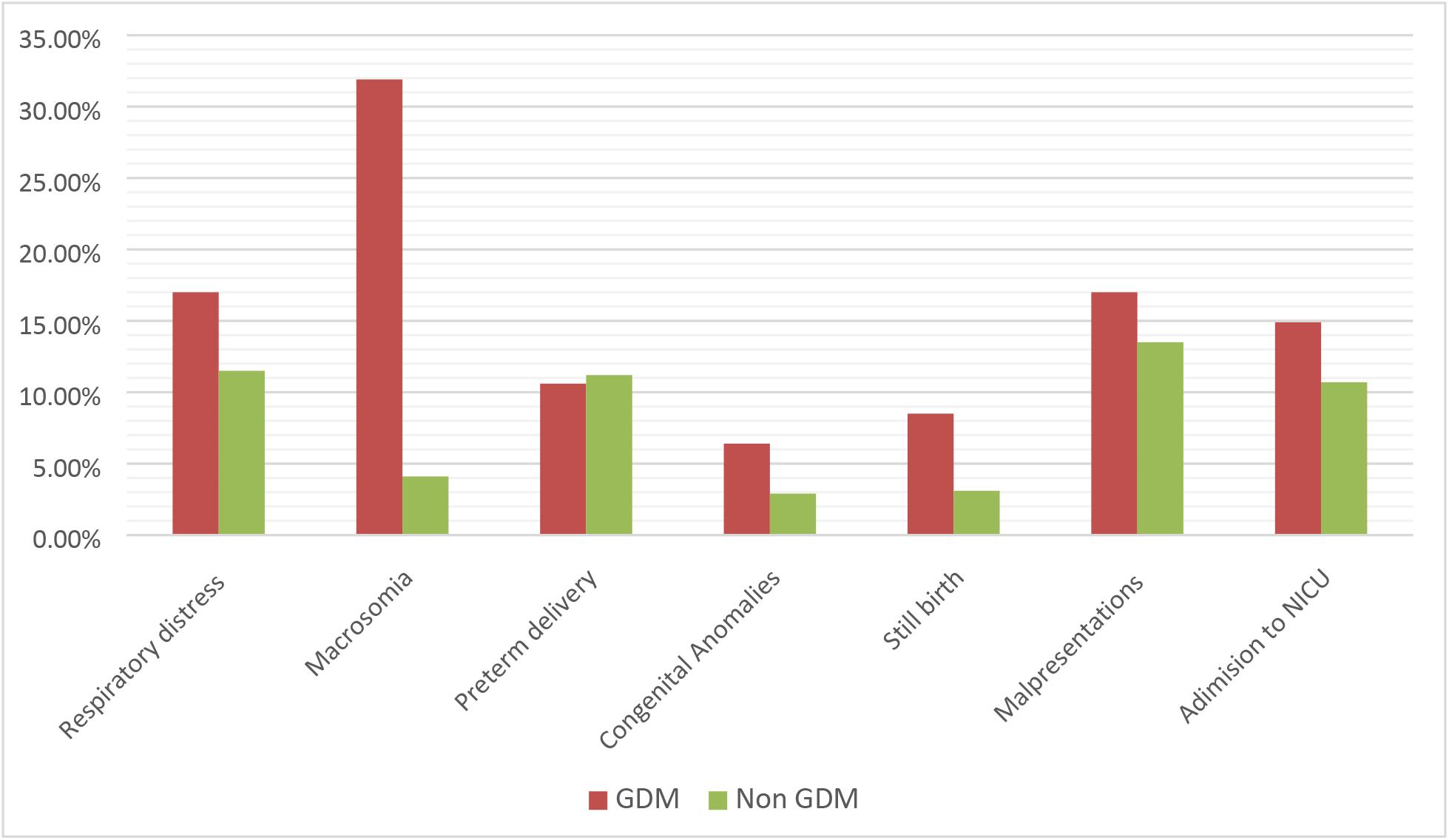
Adverse birth outcomes among mothers with and without gestational diabetes who gave birth in Hiwot Fana and Dilchora hospitals, Eastern Ethiopia, from June 2016 to April 2017.

Binary logistic regression was done to assess the association of GDM and birth outcomes. In binary logistic regression analysis Macrosomia COR=11[95% CI = 5.7-21.2] and still birth COR= 2.9[95% CI = 1.02-8.5] were significantly associated with gestational diabetes. However only macrosomia was significantly associated in multivariate analysis. Babies born to mothers with gestational diabetes were 8.5 times more likely to have macrosomia than babies born to non-diabetic mothers AOR = 8.5 [95% CI = 5.7-21.4] (Table 2).

**Table 2:**
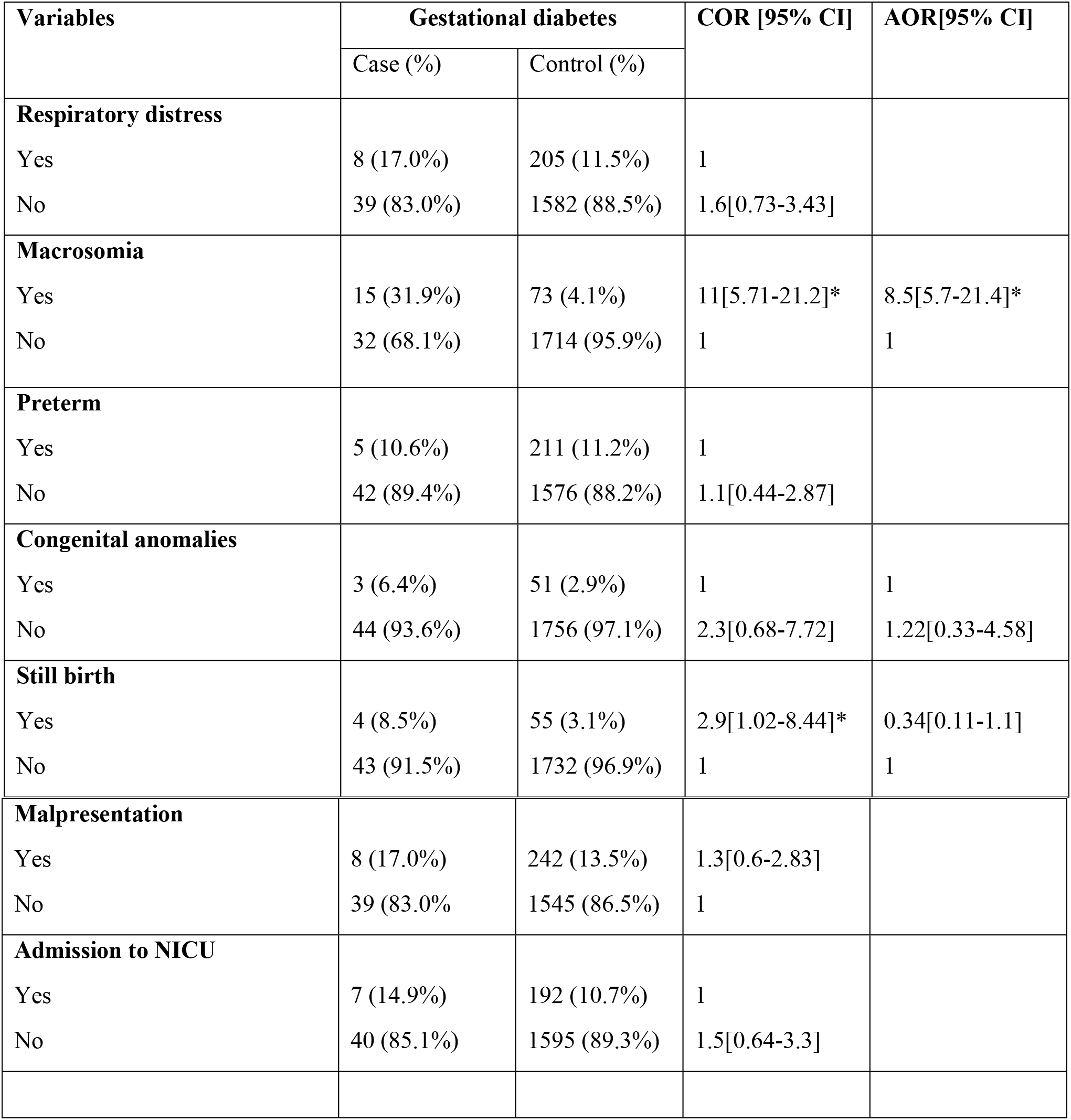
Bivariate and multivariate logistic regression analysis showing relation between adverse birth outcomes and gestational diabetes among women who gave birth in Hiwot Fana and Dilchora Hospitals from June 2016 to April 2017.

| Variables | Gestational diabetes |  | COR [95% CI] | AOR[95% CI] |
| --- | --- | --- | --- | --- |
|  | Case (%) | Control (%) |  |  |
| **Respiratory distress** |  |  |  |  |
| Yes | 8 (17.0%) | 205 (11.5%) | 1 |  |
| No | 39 (83.0%) | 1582 (88.5%) | 1.6[0.73-3.43] |  |
| **Macrosomia** |  |  |  |  |
| Yes | 15 (31.9%) | 73 (4.1%) | 11[5.71-21.2]* | 8.5[5.7-21.4]* |
| No | 32 (68.1%) | 1714 (95.9%) | 1 | 1 |
| **Preterm** |  |  |  |  |
| Yes | 5 (10.6%) | 211 (11.2%) | 1 |  |
| No | 42 (89.4%) | 1576 (88.2%) | 1.1[0.44-2.87] |  |
| **Congenital anomalies** |  |  |  |  |
| Yes | 3 (6.4%) | 51 (2.9%) | 1 | 1 |
| No | 44 (93.6%) | 1756 (97.1%) | 2.3[0.68-7.72] | 1.22[0.33-4.58] |
| **Still birth** |  |  |  |  |
| Yes | 4 (8.5%) | 55 (3.1%) | 2.9[1.02-8.44]* | 0.34[0.11-1.1] |
| No | 43 (91.5%) | 1732 (96.9%) | 1 | 1 |
| **Malpresentation** |  |  |  |  |
| Yes | 8 (17.0%) | 242 (13.5%) | 1.3[0.6-2.83] |  |
| No | 39 (83.0%) | 1545 (86.5%) | 1 |  |
| **Admission to NICU** |  |  |  |  |
| Yes | 7 (14.9%) | 192 (10.7%) | 1 |  |
| No | 40 (85.1%) | 1595 (89.3%) | 1.5[0.64-3.3] |  |

## Discussion

If not appropriately managed gestational diabetes may result in serious health complications during pregnancy, delivery and in a long term both mothers and their babies are more likely to develop type 2 diabetes. Gestational diabetes can result in higher maternal and prenatal morbidity. Different studies indicated that GDM is associated with multiple adverse birth outcomes such as; preterm delivery, still birth, respiratory distress, birth trauma and others.

In this study macrosomia was significantly associated with GDM and this finding is in line with a study done in India where macrosomia was significantly higher among women with GDM [15]. This can be explained as maternal hyperglycemia exposes the fetus to either sustained hyperglycemia or intermittent pulses of hyperglycemia and both situations prematurely stimulate fetal insulin secretion [16].

The risk of still birth is higher among pregnant women with GDM. According to the study done by Rosenstein et al the overall risk of still birth from 36 – 42 weeks was higher among a women with GDM when compared to women without gestational diabetes [17]. Similarly in this study still birth was significantly associated with gestational diabetes in binary logistic regression analysis. This can be explained as the fetus may grow slowly in uterus due to poor circulation or other conditions such as, high blood pressure or microvascular disease which can complicate diabetic pregnancy.

Preterm delivery is one of the major cause of neonatal morbidity and mortality worldwide. In Ethiopia preterm delivery is the major cause of neonatal mortality, in the year 2014 it was the cause of one third of neonatal death [18]. Some studies indicated that gestational diabetes have an association with preterm delivery, study done in Australia and Qatar revealed that preterm delivery was higher among women with GDM [19,20]. Nevertheless, other studies indicated that there is no significant association between preterm delivery and GDM. In the current study also preterm delivery was not associated with GDM.

The study done by Blanc et al showed the gestational diabetes was the independent risk factor for severe respiratory distress syndrome after 34 weeks of gestation. Respiratory distress was also significantly associated with GDM in studies conducted in Qatar and Saudi Arabia [20, 21]. However in the current study the neonatal distress was not associated with GDM. The difference might be due to the difference in the study design and number of study participants.

## Conclusion

This study revealed that macrosomia has a significant association with GDM and the odds of macrosomia was higher among mothers with gestational diabetes. Based on this study we recommend that screening for pregnant women at risk of developing GDM during the second trimester becomes mandatory to prevent macrosomia and to take appropriate interventions. Federal, regional health offices and all stake holders should work together to provide screening materials at ANC units so that the number of pregnant women screened and treated early will be screened.

## Acknowledgments

My special gratitude goes to Haramaya University for funding this research project. My deepest gratitude also goes to the data collectors and supervisor for their devotion and effective undertaking of responsibilities throughout data collection and respondents at both hospitals without whom this study would not have been realized.

## Authors’ contributions

EB - designed the study, performed the statistical analysis and drafted the manuscript. TD and KF participated in the implementation of the study and contributed to the draft manuscript. All authors have read and agreed to the final version of this manuscript and have equally contributed to its content and to the management of the case.

## ACRONYMS OR ABBREVIATIONS

ANC: Antenatal Care
BMI: Body Mass Index
DM: Diabetes Mellitus
GDM: Gestational Diabetes Mellitus
IGT: Impaired Glucose Tolerance
NICU: Neonatal Intensive Care Unit
OGTT: Oral Glucose Tolerance Test
RBS: Random Blood Sugar
WHO: World Health Organization

